# Bulked-Segregant Analysis Coupled to Whole Genome Sequencing (BSA-Seq) for Rapid Gene Cloning in Maize

**DOI:** 10.1101/357384

**Authors:** Harry Klein, Yuguo Xiao, Phillip A. Conklin, Rajanikanth Govindarajulu, Jacob A. Kelly, Michael J. Scanlon, Clinton J. Whipple, Madelaine Bartlett

## Abstract

Forward genetics remains a powerful method for revealing the genes underpinning organismal form and function, and for revealing how these genes are tied together in gene networks. In maize, forward genetics has been tremendously successful, but the size and complexity of the maize genome made identifying mutant genes an often arduous process with traditional methods. The next generation sequencing revolution has allowed for the gene cloning process to be significantly accelerated in many organisms, even when genomes are large and complex. Here, we describe a bulked-segregant analysis sequencing (BSA-Seq) protocol for cloning mutant genes in maize. Our simple strategy can be used to quickly identify a mapping interval and candidate single nucleotide polymorphisms (SNPs) from whole genome sequencing of pooled F2 individuals. We employed this strategy to identify *narrow odd dwarf* as an enhancer of *teosinte branched1,* and to identify a new allele of *defective kernel1.* Our method provides a quick, simple way to clone genes in maize.

References numbers for supporting data:figshare and NCBI SRA links to follow

## INTRODUCTION

Forward genetics remains a powerful way to ‘ask the plant’ which genes matter for a particular trait or phenotype (Mueller 2006). Because forward genetics relies on random mutagenesis, it presents an unbiased method for identifying novel genes that act in particular pathways. While forward genetic screens can reveal new genes, they can also reveal novel functions for known genes; providing a richer, more deeply nuanced view of gene function essential for a true understanding of how genes and gene networks contribute to building an organism (Nawy *et al.* 2010; Gallavotti *et al.* 2010; Vlad *et al.* 2014; Gillmor *et al.* 2016). After random mutagenesis, mutant genes are most often identified through linkage mapping.

Linkage mapping, using a combination of bulked-segregant analysis (BSA) and fine mapping, has been very successful for cloning maize genes (Gallavotti and Whipple 2015). The process starts when a mutant of interest is crossed to a wild type individual in a contrasting genetic background. The resulting F1 individuals are selfed or backcrossed, and mutants are identified in the F2 or backcross population (Fig. 1a). These mutants can be used to identify a region of increased homozygosity in a chromosomal region physically linked to the lesion causing the mutant phenotype. This region of increased homozygosity can be rapidly detected using bulked-segregant analysis (Michelmore *et al.* 1991). In bulked-segregant analysis, pools of wild type and mutant individuals are genotyped at markers spread across the genome. In chromosomal regions not linked to the mutant lesion, these markers will show a typical 1:2:1 (or 1:1 in backcross) segregation ratio. Linked markers will be homozygous for the mutant parent genotype, unless F1 recombination has happened between the mutant lesion and the marker. Once a chromosomal region has been identified using BSA, this same F1 recombination is used to place recombination breakpoints in individual mutants, and narrow the mapping interval using fine mapping (Gallavotti and Whipple 2015).

**Figure 1.**
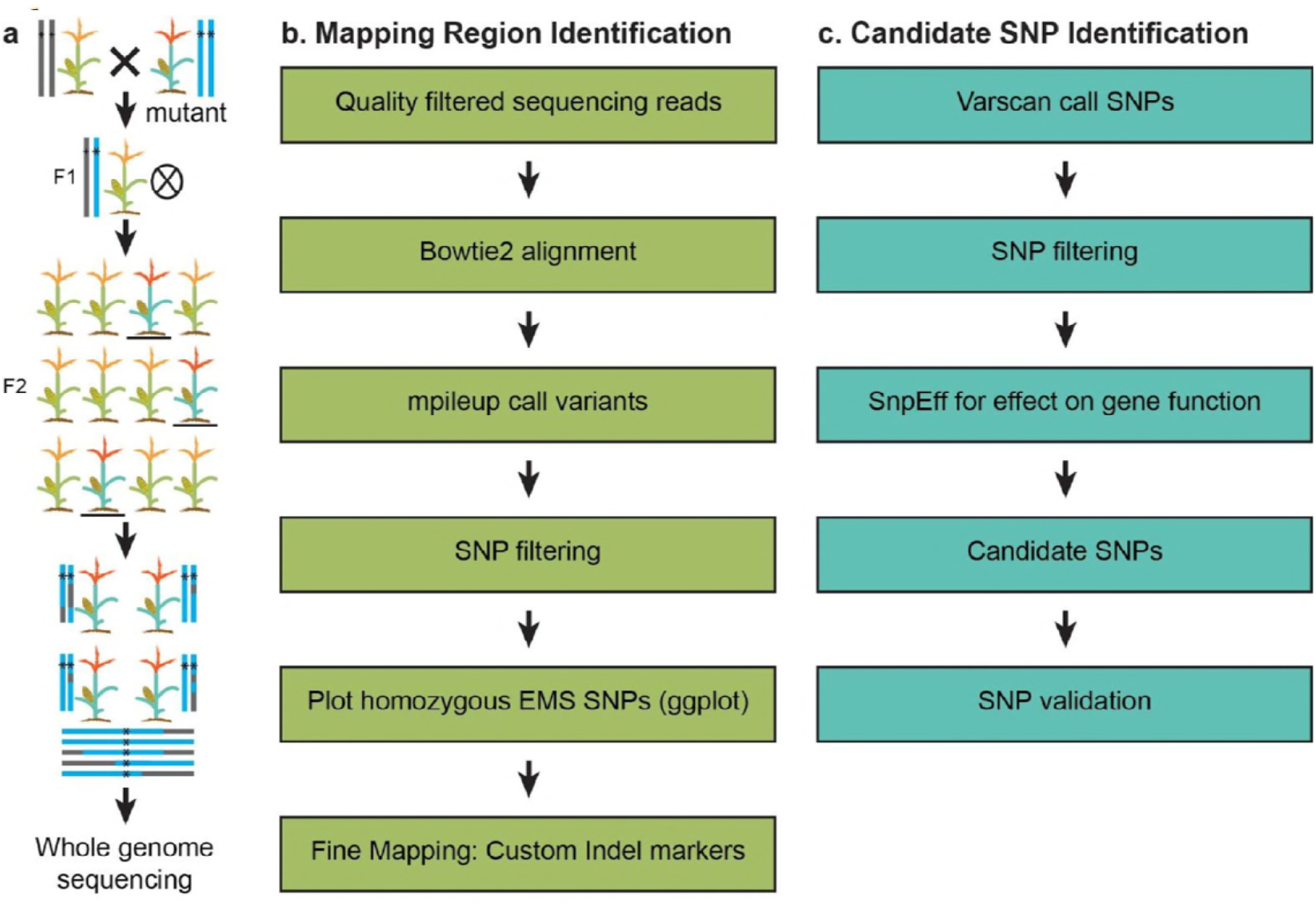
An overview of our BSA-seq pipeline. (**a**) In a BSA-Seq experiment, a plant carrying the mutant allele is crossed to or by a wild type plant of a contrasting genotype. The resulting F1 generation is selfed (or backcrossed) to generate an F2 mapping population. This mapping population is screened, and tissue from mutants pooled into a single DNA extraction. This DNA extraction is used to make a single library for whole genome shotgun sequencing. (**b**) After sequencing, we analyzed our data in Galaxy and in R to identify a chromosomal region that contains the mutant gene. (**c**) SNP filtering to identify candidate (EMS) SNPs.

The advent of next-generation sequencing (NGS) means that BSA can be used to very quickly identify a very small mapping interval, and even the causative lesion, without fine mapping. Whole genome shotgun sequencing using NGS can be used to quickly genotype mutant vs. non mutant BSA pools at many thousands of markers spread across the genome. These BSA-Seq data can reveal the genomic interval that contains the mutant gene of interest, and with enough coverage, the lesion itself (Zou *et al.* 2016). Since random mutagenesis produces lesions throughout the genome, and since NGS does not depend on established genotyping assays, BSA-Seq can be used to identify a chromosomal region using just the polymorphisms induced by mutagenesis. In this case, the contrasting genetic background used for the BSA can simply be that of an unmutagenized parent. This modification to BSA-Seq has been called MutMap (Abe *et al.* 2012). BSA-Seq has been used to identify mutant loci in *Arabidopsis thaliana*, soybean, barley, *Mimulus,* rice, sorghum, and *Brachypodium distachyon* (Schneeberger and Weigel 2011; Abe *et al.* 2012; Mascher *et al.* 2014; Woods *et al.* 2014; Ding *et al.* 2017; Song *et al.* 2017; Jiao *et al.* 2018). Although BSA-Seq has been used to identify genomic regions underlying variation in flowering time and plant height QTLs in a maize population (Haase *et al.* 2015), there have been no reports of it used to clone mutant maize genes.

Here, we report a protocol we developed for BSA-Seq in maize, and used to clone two genes recovered from EMS mutagenesis screens. We identified *narrow odd dwarf* (*nod*) as an enhancer of *teosinte branched1* (*tb1*), and identified a seedling lethal allele of *defective kernel1* (*dek1*) in a *narrow sheath (ns)* enhancer screen (Scanlon *et al.* 1996; Doebley *et al.* 1997; Lid *et al.* 2002; Becraft *et al.* 2002; Rosa *et al.* 2017). We used our *tb1* enhancer data to design insertion-deletion (indel) markers for downstream fine mapping. This fine mapping to reduce the recombination interval is critical in cases where only a single allele for a particular mutant exists, and in cases where no clear candidate lesion is identified. Our method provides a quick, easy method for cloning mutant genes in maize.

## METHODS

### Plant material and isolation of mutants

To identify enhancers and suppressors of the *tb1* (Doebley *et al.* 1997) and *ns* (Scanlon *et al.* 1996) phenotypes, we performed EMS mutagenesis screens. We mutagenized pollen homozygous for a weak allele of *tb1* in the A619 inbred (*tb1-sh*), and *ns* in an unknown background obtained from Pioneer Hi-Bred Intl. (*ns1;ns2*), using established protocols (Neuffer *et al.* 1997). After mutagenesis, M1 progeny were selfed to generate M2 populations, where we identified the mutants *tb1 enhancer* (*ten*) and *very narrow sheath* (*vns*). *ten*, which arose in the A619 genetic background, was crossed to the B73 genetic background and then selfed to generate an F2 mapping population (Fig. 1). *vns* is seedling-lethal. Therefore a wild type sibling, heterozygous at *vns*, was backcrossed to a *ns1* heterozygote fixed for *ns2 (NS1/ns1;ns2/ns2)*, and the progeny selfed to generate a mapping population. For BSA-seq, tissue samples were collected from 101 *ten* mutant individuals, and 9 *vns* mutant individuals in the mapping populations. Because the *vns* genetic background is unknown, we also collected tissue samples for 9 wild type siblings in the *vns* mapping population and for the parents of the EMS mutagenesis screen. To characterize the genetic interaction between *tb1* and *ten*, tillers were counted in families segregating *tb1-sh* and *ten*.

### DNA extractions and NGS sequencing

Tissue samples were pooled by hole punching each leaf twice to ensure equal representation of individuals. We extracted DNA from the pools of tissue using a CTAB method (Gallavotti and Whipple 2015). Libraries were prepared using the TruSeq DNA sample prep kit according to the manufacturer’s instructions (Illumina). We sequenced the *ten* mutant pool on an Illumina Hi-Seq 2500 at Brigham Young University. Sequenced reads were 125bp long with paired-ends. We sequenced *vns* mutants, *vns* wild type siblings, and unmutagenized parents on an Illumina Hi-Seq 2500 at Cornell University. Sequenced reads from *vns* were 150bp long with paired ends.

### NGS quality control and read alignment

We used the genomic tools hosted on Galaxy web portal to process our Next-Gen sequencing data (Afgan *et al.* 2016). An overview of our process is shown in Figure 1. We used FastQC (v. 0.69) to determine the quality of our sequencing data, and established a PHRED quality cutoff of 20 based on this analysis (Ewing *et al.* 1998; Andrews 2014). We used FASTQ Groomer (v. 1.1.1) to convert our FASTQ files to Sanger format for input into downstream Galaxy tools (Blankenberg *et al.* 2010). Trimmomatic (v. 0.36.3) was used for sliding window trimming averaged across 4 base pairs with average quality >20 and for adapter sequence removal (Bolger *et al.* 2014). The Galaxy tool cat, which concatenates datasets tail-to-head (cat) (v. 0.1.0) was used to join sequencing files together if data was split from multiple Illumina sequencing runs (Gruening 2014). We used Bowtie2 (v. 2.3.2.2) to align sequencing reads to the B73 reference genome version 4, release 56 and to generate an index (Langmead and Salzberg 2012; Jiao *et al.* 2017). Read alignment was assessed with SAMtools Flagstat (v. 2.0) (Li *et al.* 2009).

### Variant calling

SAMtools mpileup (v. 2.1.3) was used to output variants from the indexed bam file into pileup or vcf files (Li *et al.* 2009). Pileup files were generated for identification of the mapping region and vcf files were generated for identification of candidate SNPs. To filter variant files for mapping region identification, we used SAMtools filter pileup (v. 1.0.2), a custom Galaxy tool, to filter based on read coverage (Li *et al.* 2009). To generate a vcf file for candidate SNP identification, we used Varscan (v. 0.1). We used a minimum homozygous calling frequency of 0.99, and filtered out SNPs with low read coverage (coverage <8) (Koboldt *et al.* 2012). Varscan (v. 0.1) was also used to call SNPs in other A619 mutant sequence data and to call SNPs in the *vns* wild type siblings and unmutagenized parents.

### Data processing for plotting and SNP filtering

The filtered pileup file was split into 10 chromosomes. At each nucleotide position in the pileup file, we calculated variant allele frequency by dividing the number of reads that differed from the B73 reference sequence by the quality adjusted total number of reads at that position. This variant allele frequency was added as a new column in the pileup file. Nucleotide position, reference allele, alternate allele, coverage, and variant allele frequency were extracted from the pileup file to create a variant file for downstream analysis in R (Wickham and Francois 2015; Wickham 2016). Similarly, SNP position files were produced for *vns* wild type siblings and unmutagenized parents, and A619 SNPs. The A619 SNPs we used were common to 5 mutant sequencing datasets (including *ten*) and were generated using the R package findCommonVariants (Shi *et al.* 2013) (File S1).

### Mapping region identification

Once we had a set of homozygous EMS SNPs particular to each of our mutants, we plotted these data against the B73 reference genome, to identify chromosomal regions enriched for homozygous EMS SNPs, presumably in linkage with the causative lesions. Then we filtered positions with coverage >100 to remove highly enriched positions in our data. To reduce noise that might come from mapping or sequencing error, we only retained SNPs that had read coverage within one standard deviation of the mean (calculated once enriched positions had been removed). For *vns*, we removed SNPs that were in the wild type sibling and unmutagenized parent datasets. We used ggplot2 to plot the number of homozygous variant positions that differed from the B73 reference genome, per 1Mbp chromosomal bin (Wickham and Francois 2015; Wickham 2016). For these analyses, nucleotide positions that differed from the reference genome at a frequency greater than or equal to 0.99 were defined as homozygous variants.

### SNP filtering for candidate SNP identification

Once we had identified likely mapping intervals for *ten* and *vns*, we searched for potentially causative lesions in each of these intervals. Here, we returned to the vcf file generated using Varscan, and removed background SNPs (A619, wild type sibling, and/or parental SNPs) as before. We reasoned that it was highly unlikely that the causative SNPs could be any known SNP present in any maize inbred. Therefore, we applied an additional SNP filtering step, and removed all the maize Hapmap 3.2.1 SNPs (Merchant *et al.* 2016; Bukowski *et al.* 2018). Once we had a set of positions enriched for SNPs unique to each of our particular mutant pools, we filtered out SNPs that were not homozygous and were not canonical EMS changes (G to A or C to T) (Till *et al.* 2004). The final set of homozygous EMS SNPs unique to each dataset was the input for an analysis of likely SNP effects on gene function.

To identify SNPs in our filtered datasets that might negatively affect gene function, we used SnpEff (version 4.3a) (Cingolani *et al.* 2012). SnpEff identifies nonsense SNPs and splice site mutations as likely to have highly deleterious effects on gene function, and all missense SNPs as likely to have moderate effects on gene function (Cingolani *et al.* 2012). Functional annotations for the genes disrupted by likely moderate- and high-effect candidate SNPs were obtained from Gramene (Tello-Ruiz *et al.* 2018). Candidate SNPs were validated through Sanger sequencing and complementation crosses.

### Indel marker design and fine mapping

We reasoned that in cases when there are more than one gene candidate, or when there are no clear candidates, reducing the mapping interval using fine mapping would be very useful. We further reasoned that the NGS data from a conventional BSA-Seq experiment could be used to quickly design many custom indel markers. As a proof of concept, we designed custom markers to refine our *ten* mapping interval, using the individual mutant leaf samples. We used Varscan (v. 0.1) to identify insertion and deletion polymorphisms (indels) in our mapping interval that differentiated our mutant pool from the B73 reference genome. We chose indels that were 15 base pairs in length or longer, and designed primers to flank these indels by 100-150bp (Table S1). DNA was extracted from each of the 101 F2 mutants originally pooled into one NGS library and used for Indel PCR (Gallavotti and Whipple 2015).

Size differences between the resulting PCR products were detected on 3.5% agarose gels.

### Data Availability

Supplemental files are available at figshare. File S1 contains A619 SNP positions. Raw sequencing data available at NCBI SRA (link to follow). Code for analyses and figure generation is available at the Bartlett lab’s github (link to follow).

## RESULTS

### Mutant phenotypes

From an EMS mutagenesis screen of *tb1-sh* mutants in the A619 genetic background, we recovered a dwarf mutant with additional tillers that we called *tb1 enhancer* (*ten*) (Fig 2a). Similarly, *very narrow sheath* (*vns*) was uncovered in an EMS mutagenesis screen looking for enhancers of the *narrow sheath* (*ns*) phenotype. *vns* single mutants are seedling lethal and often fail to develop more than 3 leaves (Fig. 2c).

**Figure 2.**
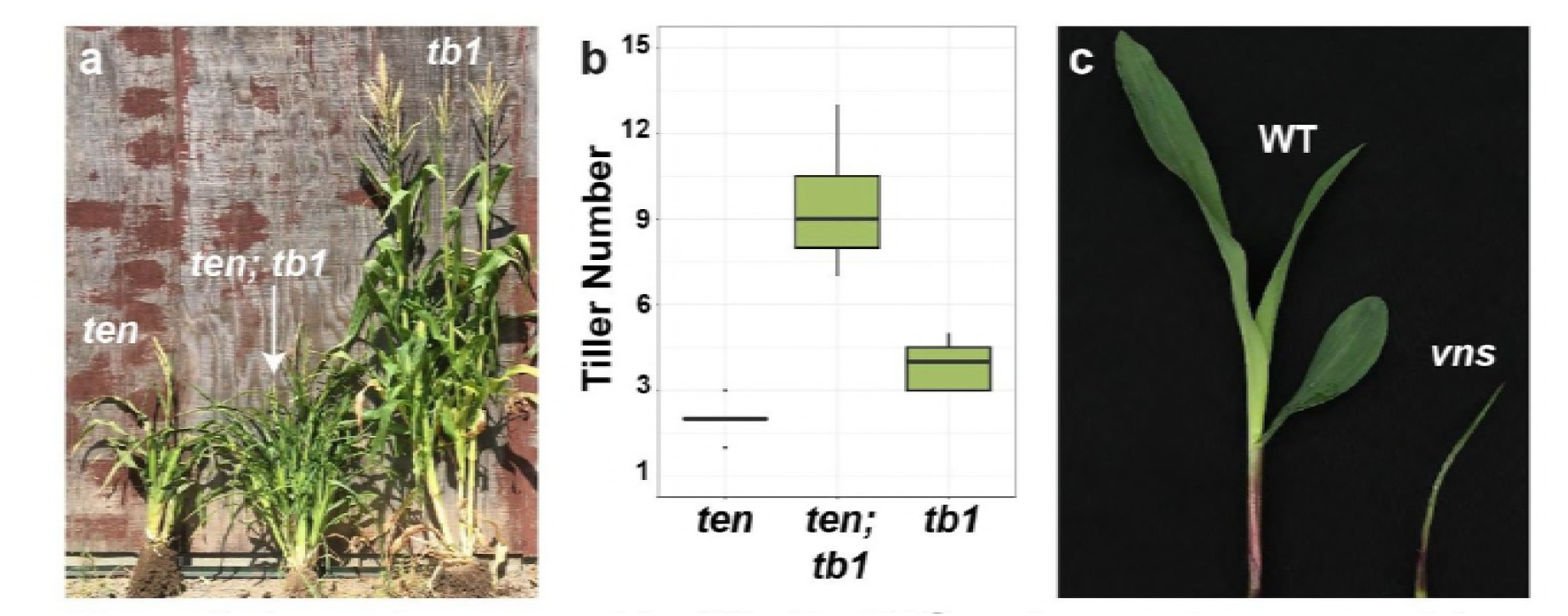
*ten* and *vns* were identified in EMS mutagenesis screens. (**a**) *ten* (left), *tb1-sh ten* (middle), *tb1-sh* (right). (**b**) Quantification of tillers in a 9:3:3:1 population of *entb1* and *tb1-sh* shows a synergistic interaction between *entb1* and *tb1-sh*. (**c**) Wild type (left) and *vns* mutant (right) exhibiting reduced growth at the seedling stage.

To determine the genetic interaction between *ten* and *tb1,* we counted tillers in a population segregating both *tb1-sh* and *ten*. Tiller number (per plant) was counted in 15 *tb1-sh* single mutants, 15 *ten* single mutants and 15 *tb1-sh*; *ten* double mutants. *tb1-sh* and *ten* single mutants had an average of 4 and 2 tillers respectively. *ten tb1-sh* double mutants had an average of 9 tillers, showing a synergistic interaction between *ten* and *tb1-sh* in tiller development (Fig. 2b).

### Both *ten* and *vns* are on chromosome one

To identify the causative lesions underlying the *ten* and *vns* mutant phenotypes, we sequenced pooled DNA from 101 *ten* individuals and 9 *vns* individuals. We sequenced *ten* pools and *vns* pools on an Illumina platform with outputs of 125 bp and 150bp paired end reads, respectively. For *ten*, we recovered 450,665,452 reads, and for *vns* we recovered 311,711,046 reads. Using Bowtie2, we mapped our reads to version 4 of the maize B73 reference genome (Jiao *et al.* 2017). We analyzed genome mapping and coverage using SAMtools flagstat (Li *et al.* 2009). For *ten*, 97% of our reads (438,746,898) mapped and 91% were properly paired (409,548,662). For *vns*, 90% of our reads (279,239,524) mapped and 77.5% were properly paired (241,580,812). Coverage of all *ten* and *vns* mapped reads was 26-fold and 17-fold respectively (Table 1). The combined dataset that we used to generate the A619 SNP dataset included 1.1 billion mapped reads, for a mean coverage of 59X (File S1). We called SNPs from our Bowtie2 alignment with SAMtools mpileup and used the Galaxy filter pileup plugin to filter out SNPs with coverage <8 (Li *et al.* 2009).

**Table 1.** Sequencing data summary for *ten* and *vns*

| Measure | *ten* | *vns* |
| --- | --- | --- |
| Reads | 450,665,452 | 311,711,046 |
| Mapped reads | 438,746,898 | 279,239,524 |
| % Mapped Reads | 97.36 | 89.58 |
| Paired reads | 409,548,662 | 241,580,812 |
| Coverage | 26.06 | 16.59 |

To identify the the chromosomal regions corresponding to *ten* and *vns*, we searched for regions of the genome enriched for homozygous variants in the mutant pools. Before searching for this enriched homozygosity, we removed high coverage SNPs (>100) from both datasets, and retained only those SNPs for each chromosome that had read coverage within one standard deviation of the mean (Table 1, Table S2). *ten* was recovered from a mutagenesis experiment in the A619 genetic background, and then crossed to B73 to generate the F1. Therefore, we were looking for a genomic region enriched for homozygous variants that differed from the B73 reference genome. To identify this region, we plotted the number of homozygous SNPs that differed from B73 (variant frequency >=0.99), per 1Mbp bin. Plotting homozygous variants in the *ten* dataset quickly identified a peak on the top of chromosome 1, between 0 and 20Mbp (Fig. 3a). Plotting only homozygous canonical EMS SNPs (G to A and C to T transitions) also identified a peak on chromosome 1, although it was smaller. We also identified several much smaller peaks at the telomeres of chromosome 1, 2, and 5 that contained regions with protein coding genes. These telomeric peaks are likely caused by differences between our B73 stocks, and those that were used to generate the B73 reference genome, either because of their histories in our own labs, or because of their provenance (Liang and Schnable 2016). We focused on the 0-20Mbp peak on chromosome 1 as the *ten* mapping interval because it was much bigger than any of the telomeric peaks. In addition, the peak at the top of chromosome 1 had broad shoulders, distinct from the sharp peaks we saw at the telomeres of chromosomes 1, 2, and 5. The 0-20Mbp peak on chromosome 1 is what we expected from a segregating locus mapped by BSA, where each genome in the pool will have different recombination breakpoints surrounding the mutant lesion. The *ten* mapping interval defined by this initial analysis included 623 genes.

**Figure 3.**
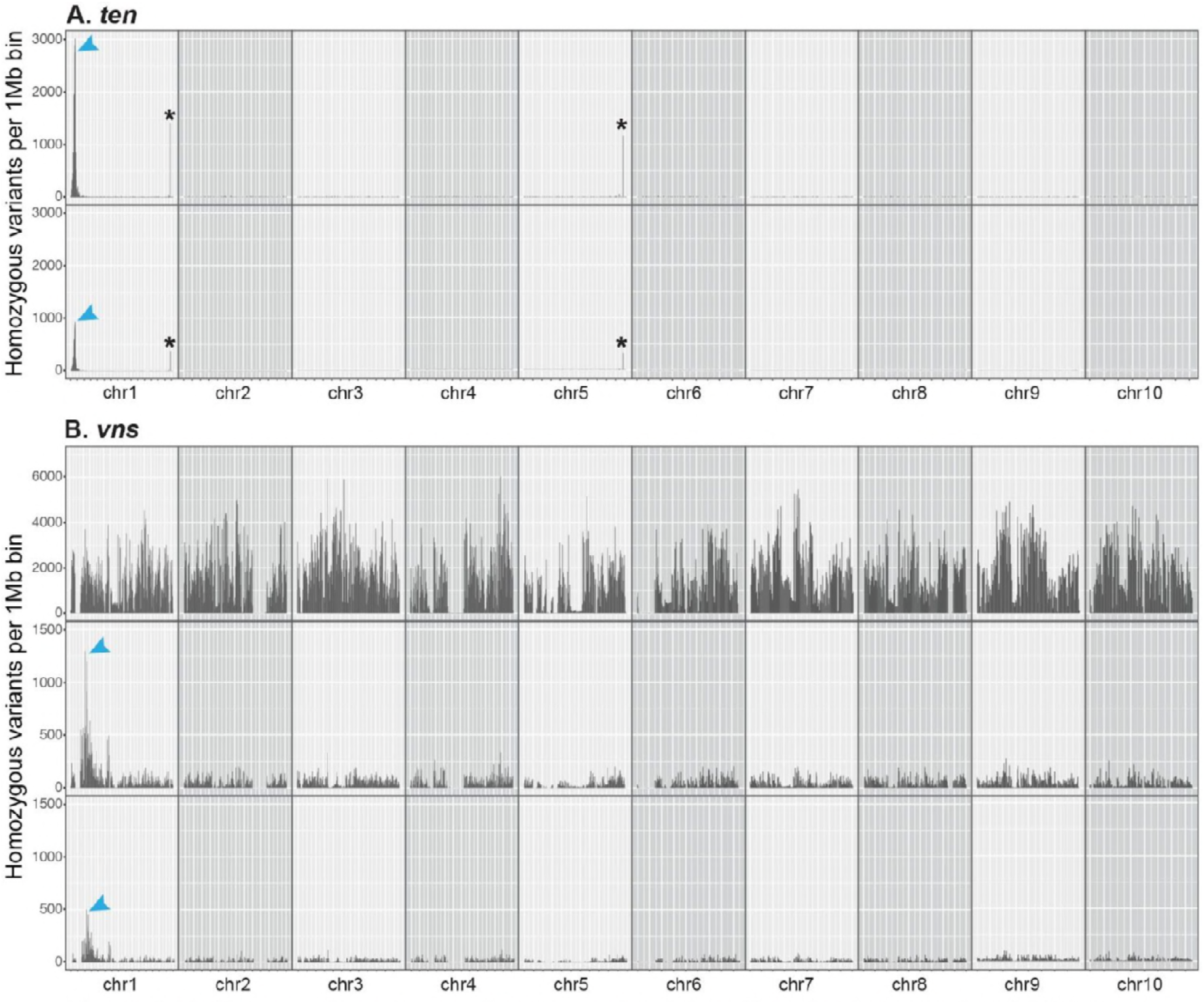
Both *ten* and *vns* are on chromosome 1. (**a**) *ten* likely lies between 0 and 20Mbp on chromosome 1. Plotting the number of homozygous positions that differ from the B73 reference genome (per 1Mbp chromosomal bin) reveals a large peak on chromosome 1 (top). This peak is still visible, but smaller, if only homozygous EMS SNPs are plotted (bottom). (**b**) *vns* likely lies between 40 and 70Mbp on chromosome 1. Plotting the number of homozygous positions that differ from the B73 reference genome (per 1Mbp chromosomal bin) reveals no distinct peaks (top). After filtering out parental, wild type sibling, Hapmap 3.2.1 and background SNPs, a large peak on the short arm of chromosome 1 is revealed (middle). This peak is still visible, but smaller, if only homozygous EMS SNPs are plotted (bottom).

For *vns*, identifying a chromosomal location was not quite as simple because *vns* was identified in an unknown genetic background, and a *vns* heterozygote was crossed to an unmutagenized parent, and not outcrossed to any reference genotype (e.g. B73). Thus, plotting only homozygous *vns* variants revealed, as expected, extensive homozygosity distinct from the B73 reference across the genome (Fig. 3b). There was still no clear peak when we plotted only canonical EMS SNPs. Therefore, to determine background SNPs that could be removed from our *vns* dataset, we sequenced separate pools of both unmutagenized parents and wild type siblings alongside the *vns* mutant pool. After removing these parental and wild type sibling SNPs, as well as all the HapMap 3.2.1 and A619 SNPs, we found a region of high homozygosity on the short arm of chromosome 1 between 30 and 70Mbp. The *vns* interval was larger compared to the *ten* region, perhaps due to the small number of individuals (9) included in each *vns* pool, or the increased recombination at the end of chr.1 where *ten* is located. As with *ten*, plotting only EMS SNPs still revealed a (smaller) peak. The presence of many non-EMS SNPs in the *vns* mapping interval is likely a result of heterozygosity in the parental *ns1 ns2* stock. The final *vns* mapping interval included 651 genes.

### Fine mapping of *ten* using custom indel markers quickly reduced the mapping interval

Both the *ten* and *vns* pools were sequenced fairly deeply (Table 1), and our sequencing data was thus likely to contain both causative lesions. In a case where mutant pools might be sequenced to a shallower depth, and the causative lesion might be missed, we reasoned that the sequencing data could be used to design markers for fine mapping to reduce the mapping interval. This fine mapping will also be critical for reducing the mapping interval when only a single mutant allele is available, or if there are multiple gene candidates in the mapping interval. As a proof of concept, we used our NGS data to design five insertion/deletion (indel) markers to refine the *ten* interval (Gallavotti and Whipple 2015) (Table S1). We used these markers to genotype 101 *ten* individuals from the F2 mapping population and narrowed the *ten* interval to a region that includes 74 genes between 11.1 and 13.1 Mbp (Fig. 4a). Thus, we were able to quickly reduce the *ten* mapping interval and simplify the evaluation of candidate SNPs.

**Figure 4.**
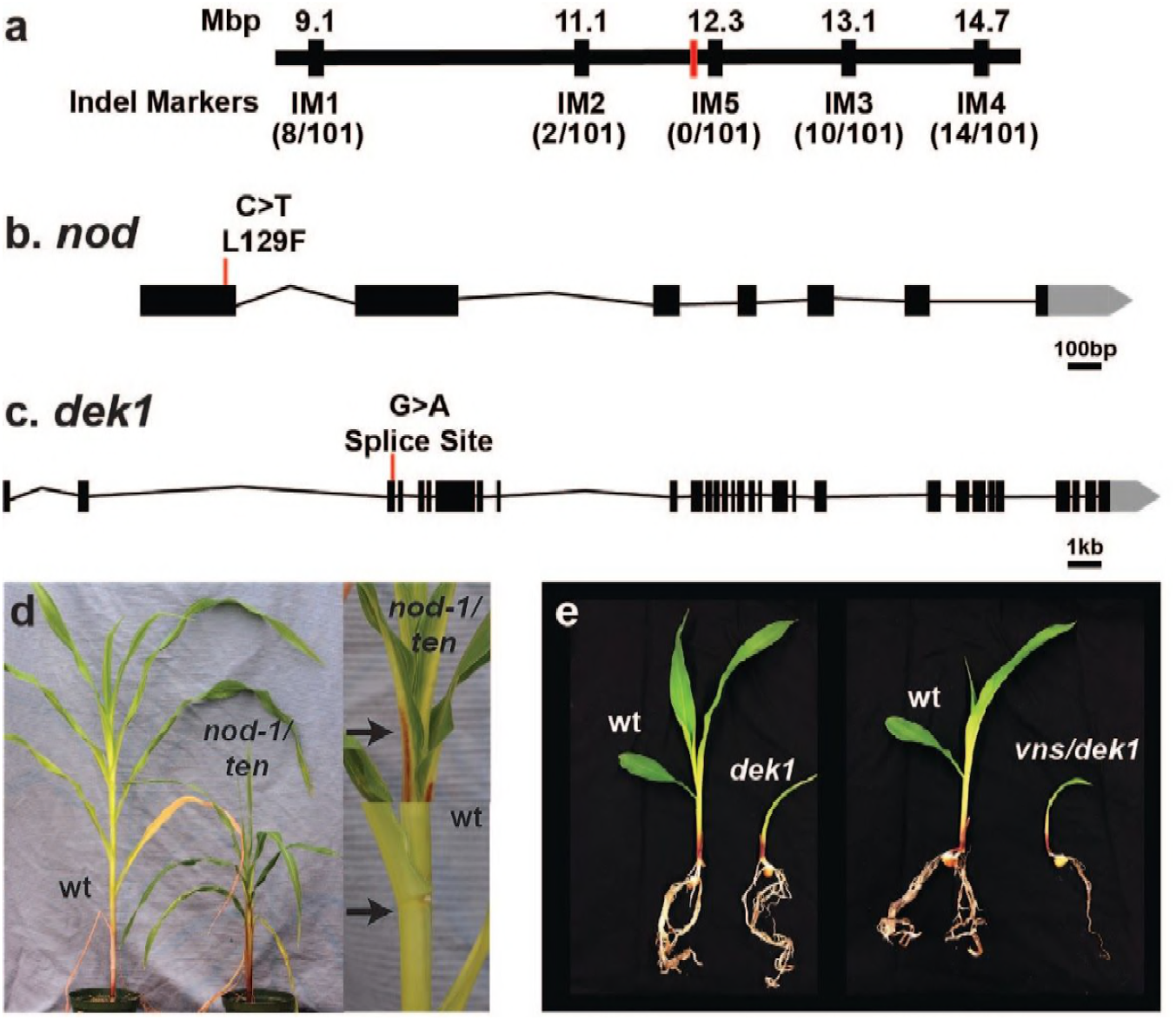
*ten* and *vns* encode alleles of *nod* and *dek1,* respectively. (**a**) Fine mapping of *ten* using custom indel markers reduced the mapping interval to a region containing 74 genes. The approximate position of *nod-ten* is marked in red. (**b**) The *ten* lesion in *nod* is a missense mutation. (**c**) The *vns* lesion in *dek1* is a splice site mutation. (**d**) *ten* fails to complement *nod.* As in *nod-1* mutants (Rosa *et al.* 2017), the ligule and auricle are both absent in the leaves of *ten/nod-1* F1 plants (top). The black arrow indicates the ligule in a wild type plant (bottom), and the lack of a ligule in a *ten/nod-1* F1 plant. (**e**) *vns* fails to complement *dek1*.

### SNP filtering and candidate SNP identification

Next, we wanted to identify candidate SNPs in our BSA-seq mapping regions. We used a variant calling program, Varscan, to call genotypes at variant positions based on allele frequency (Koboldt *et al.* 2012). Although we could have investigated the filtered EMS SNPs from our pileup file used to determine the mapping region directly, we chose to use Varscan because it outputs a variant call format (vcf) file, needed for downstream candidate SNP analyses (SnpEff) (Fig. 1c).

Varscan identified 36,186,873 and 36,790,695 SNPs genome-wide in the *ten* and *vns* datasets, respectively (minimum read depth of 8) (Table 2). We removed putative background SNPs from these datasets, starting with all the known Hapmap 3.2.1 variants (Bukowski *et al.* 2018), all A619 variants (File S1), and SNPs from wild type siblings and unmutagenized parents (*vns*). We next filtered out all but canonical EMS SNPs (G to A or C to T) (Till *et al.* 2004). A lesion leading to a recessive mutant phenotype should be homozygous, so we also filtered out non-homozygous variants. Sequencing error or pool contamination with wild type individuals could both lead to the causative SNP being erroneously filtered out as heterozygous. Similarly, if the causative lesion is a non-canonical EMS mutation, it could also be prematurely removed from the list of candidate SNPs. Therefore, we applied these filters last. After SNP filtering, we were left with 50 homozygous EMS SNPs in the *ten* mapping interval, and 427 in the *vns* interval (allele frequency>0.99).

**Table 2.** Varscan SNP data summary for *ten* and *vns*(coverage>8).

| SNPs | *ten*<br>(ROI=0-20Mbp) | *vns*<br>(ROI=40-70Mbp) |
| --- | --- | --- |
| <u>Genome-Wide:</u> |  |  |
| Total SNPs (genome wide) | 36,186,873 | 36,790,695 |
| Homozygous EMS SNPs after filtering | 148 | 7,899 |
| <u>Region of Interest (ROI)</u> |  |  |
| Total SNPs | 266,837 | 470,763 |
| Homozygous EMS SNPs after filtering | 50 | 427 |
| High effect homozygous EMS SNPs | 0 | 1 |
| Moderate effect homozygous EMS SNPs | 4 | 2 |

To identify which SNPs in these final sets might have deleterious effects on gene function, we ran these SNPs through SnpEff (Cingolani *et al.* 2012). SnpEff calls any putative missense SNPs as likely moderate effect SNPs. Likely high effect SNPs are those that are splice site variants or introduce an early stop codon. Inversions, insertions, and deletions in exons are also identified as likely high effect SNPs. SnpEff identified 0 SNPs of high effect, and 4 SNPs of moderate effect in our *ten* mapping interval. The *vns* mapping interval included 1 SNP of likely high effect, and 2 SNPs of likely moderate effect (Table 3).

**Table 3.** Candidate SNPs for *ten* and *vns*

| Chr | Position | Ref | Alt | Gene ID | SnpEff Call | Effect | Functional Annotation |
| --- | --- | --- | --- | --- | --- | --- | --- |
| *ten* |  |  |  |  |  |  |  |
| 1 | 12172055 | C | T | Zm00001d027722 | Moderate | L129F | *narrow odd dwarf (nod)* |
| 1 | 12286537 | C | T | Zm00001d027725 | Moderate | P208L | 3-deoxy-D-arabino-heptulosonate 7-phosphate synthase 1 (DHS1) |
| 1 | 12631437 | C | T | Zm00001d027752 | Moderate | G139E | SH2 Domain Containing Protein |
| 1 | 13337447 | G | A | Zm00001d027775 | Moderate | D67N | Histone H2A 8 |
| *vns* |  |  |  |  |  |  |  |
| 1 | 47570165 | G | A | Zm00001d028818 | High | Splice Site | *defective kernel1 (dek1)* |
| 1 | 53173992 | G | A | Zm00001d028965 | Moderate | A38V | Protein of unknown function |
| 1 | 53174101 | G | A | Zm00001d028965 | Moderate | R2C | Protein of unknown function |

### Candidate SNP validation

In our *ten* dataset there were four remaining moderate-effect SNPs: in genes encoding a 3-deoxy-D-arabino-heptulosonate 7-phosphate synthase 1 (*DHS1*) (Zm00001d027725), an SH2 domain protein (Zm00001d027752), Histone H2A 8 (Zm00001d027775), and in a known maize gene - *narrow odd dwarf* (*nod*) (Zm00001d027722). The phenotype of *nod* mutants strongly resembled *ten* single mutants (Rosa *et al.* 2017). Therefore, we Sanger sequenced *nod* in one of our *ten* mutants, and confirmed the missense mutation identified through NGS (Fig 4b). In addition, we crossed *ten* to heterozygous *nod* plants and examined the F1 progeny. We found that 12 of 26 F1 plants were short, and lacked ligules (Fig 4d), as is the case with *nod* mutants (Rosa *et al.* 2017). Therefore, we concluded that *ten* is an allele of *nod,* and henceforth will refer to it as *nod-ten.*

In our *vns* dataset, there were 2 likely moderate-effect SNPs and 1 high effect SNP. The high effect SNP was in a donor splice site of a known maize gene, *defective kernel1* (*dek1*) (Zm00001d028818) (Fig. 4c). *dek1* mutants contain a root primordium but lack a shoot structure (Neuffer *et al.* 1997), similar to what was observed in *vns* mutants. To determine if the SNP in *dek1* was responsible for the *vns* phenotype, we crossed heterozygous *vns* plants with heterozygous *dek1* plants. In the F1 progeny, 9 of 36 plants were very small, and died shortly after producing only a small number of leaves. Similarly, 7 of 36 F1 progeny of the heterozygous *dek1* self exhibited the same phenotype. Thus, *vns* failed to complement the *dek1* mutant phenotype, indicating that *vns* is an allele of *dek1.* Henceforth we will refer to *vns* as *dek1-vns*.

## DISCUSSION

We developed a BSA-Seq method for cloning genes in maize. Using this method, we identified and validated causative SNPs for two separate EMS-induced mutants. We have provided a protocol for conducting these analyses using Galaxy, as well as all of our annotated code. In addition, we generated a SNP variant file for the A619 genetic background (File S1). This has the potential to be useful for anyone working in A619 - including for making a synthetic A619 reference genome (McKenna *et al.* 2010).

To minimize the size of the chromosomal region recovered using BSA-Seq, a number of factors need to be considered. Factors that influence the precision of BSA-Seq include the size of the mutant pool sequenced, sequencing depth, and background levels of recombination across the genome. In the case of *nod-ten*, we sequenced a fairly large pool (101 mutant individuals), and *nod* is in a region of the genome where recombination is high (Gore *et al.* 2009). Therefore, the *nod-ten* peak is quite steep and tight, encompassing ~20Mbp on the long arm of chromosome 1. In the case of *dek1-vns*, the peak is broader, encompassing ~30Mbp on the long arm of chromosome 1. Peak size here is likely influenced by both the small pool size used for sequencing (9 mutant individuals), and by lower recombination rates in the *dek1* chromosomal region. Mean coverage in both datasets was also fairly high, allowing for both causative lesions to be captured. To identify just the genomic interval that contains a particular gene, sequencing depth could be reduced, but the chances of catching the causative lesion will be similarly reduced. If no clear candidate lesion is recovered, and/or there are many genes in a BSA-Seq mapping interval (even the fairly tight *nod-ten* BSA interval included 623 genes), fine mapping would become essential. In addition, the chromosomal region, and therefore the recombination rates at a locus, cannot be predicted before starting a BSA-Seq experiment. Thus, we recommend a big pool of mutants (100-200), sequenced to the deepest level affordable, for a successful BSA-Seq experiment.

For *dek1-vns*, we used a modification of BSA-Seq that has been called MutMap (Abe *et al.* 2012). In MutMap, instead of generating an F1 between contrasting genotypes, the F1 comes from a mutant backcrossed to a wildtype, unmutagenized individual in the same genetic background as the mutant. Thus, MutMap introduces no additional genetic variation in the initial F1 cross. Instead, MutMap relies on co-segregation of induced SNPs with the mutant phenotype, allowing for the identification of a region of increased homozygosity in the mutant pool. MutMap would likely also have been successful in identifying *nod-ten.* In both of our datasets, just mapping the EMS mutations reveals clear peaks corresponding to the mutant genes (Fig. 3). One advantage that MutMap offers is that no additional genetic variation is introduced in the F1 cross. This genetic variation can suppress or enhance mutant phenotypes, which can make scoring F2 populations challenging. However, maize geneticists are actively working to identify the modifiers in contrasting genetic backgrounds (Bolduc *et al.* 2014). In these cases, conventional BSA-Seq could be used to identify both causative lesions and modifiers in one step (Song *et al.* 2017).

The most important advantage of BSA-Seq is its simplicity, both in terms of sample collection and data analysis. While BSA-Seq is used extensively in other taxa (Schneeberger and Weigel 2011; Abe *et al.* 2012; Mascher *et al.* 2014; Woods *et al.* 2014; Ding *et al.* 2017; Song *et al.* 2017; Jiao *et al.* 2018), gene mapping via Bulked-Segregant RNA-Seq (BSR-Seq) is more often used for cloning maize genes (Liu *et al.* 2012; Li *et al.* 2013; Nestler *et al.* 2014; Tang *et al.* 2014). In BSR-Seq, RNA from a pool of mutants and RNA from a pool of non-mutants is used to make RNA-Seq libraries (Liu *et al.* 2012). These libraries are sequenced, and the resulting reads mapped to the maize reference genome. BSR-Seq offers a very good method for genome reduction, thus increasing sequencing depth without increasing cost. If the RNA-Seq is performed using the right tissue at the right developmental stage, BSR-Seq offers the advantage of differential gene expression data as well as mapping information. In contrast, since it relies on DNA extraction, BSA-Seq sample collection can be done at any developmental stage, and from any tissue. Although bulk RNA extractions are not extraordinarily challenging, DNA extractions are far simpler, and can be performed by relatively inexperienced trainees in the lab. This technical simplicity offers the advantage of being able to involve high school students and junior undergraduates in authentic research experiences (Lopatto *et al.* 2014). In addition, the data analysis for BSR-Seq is not as straightforward as it is for BSA-Seq. In BSR-Seq, allele-specific expression must be accounted for, as well as differential expression of genes not linked to the mutant gene in mutant vs. wild-type pools (Liu *et al.* 2012). Here, too, simplicity offers speed for the experienced researcher, and excellent training opportunities for beginning scientists.

Another critical advantage of BSA-Seq is that even if the lesion is not captured, a researcher is immediately poised to design indel markers and commence fine-mapping in an identified genomic interval. This has the potential to be especially useful when a candidate lesion is not clear. This is likely to be more problematic when the mutant under study is not from an EMS mutagenesis experiment. In cases such as these, which are also fairly common in maize (Vollbrecht *et al.* 1991; Thompson *et al.* 2009; Whipple *et al.* 2011), high homozygosity that is polymorphic with a reference genome will still reveal the chromosomal location of the causative lesion and streamline subsequent fine mapping.

Technical and conceptual advances will eliminate some, but not all, sources of uncertainty when it comes to capturing the candidate lesions underlying mutant phenotypes. With shallow read depths, causative lesions may not be captured; but deep coverage is likely to become ever more attainable as NGS costs drop, thus eliminating this source of uncertainty. There may be a number of potentially causative lesions in a candidate region; but as the maize pan-genome is resolved (Hirsch *et al.* 2014; Bukowski *et al.* 2018), the pool of background SNPs that can be eliminated gets ever-deeper when cloning clear null mutations unlikely to be present in natural variation. Other potential sources of uncertainty will likely never go away. For example, filtering out all of the HapMap and pan-genome SNPs will be less useful when trying to identify natural modifiers. BSA-Seq and fine mapping will be particularly helpful in narrowing down a candidate region in this case; and also when a causative lesion is not in the coding sequence of an annotated gene; or when mutant pools are contaminated with wild type or heterozygous samples. Maize geneticists often face two particular challenges: (1) only one mutant allele might be available when an experiment starts, which makes candidate validation challenging; and (2) lesions are not necessarily of a defined type (e.g. G to A or C to T transitions), which makes SNP filtering challenging. Thus, BSA-Seq has advantages that are likely to prove useful for many maize mutants.

## Acknowledgements

The authors thank Courtney Babbitt and Beth Thompson for helpful discussion; Sarah Hake for the kind gift of the *tb1-sh* allele; and Ed Wilcox at the BYU DNA sequencing facility for technical assistance. This work was supported by a USDA-Hatch grant to PI Bartlett (MAS00501), the Biology Department at the University of Massachusetts Amherst, and an NSF PGRP grant to PI Whipple (IOS-1339332).

